# Quantifying differences between passive and task-evoked intrinsic functional connectivity in a large-scale brain simulation

**DOI:** 10.1101/250894

**Authors:** Antonio Ulloa, Barry Horwitz

## Abstract

Establishing a connection between intrinsic and task-evoked brain activity is critical because it would provide a way to map task-related brain regions in patients unable to comply with such tasks. A crucial question within this realm is to what extent the execution of a cognitive task affects the intrinsic activity of brain regions not involved in the task. Computational models can be useful to answer this question because they allow us to distinguish task from non-task neural elements while giving us the effects of task execution on non-task regions of interest at the neuroimaging level. The quantification of those effects in a computational model would represent a step towards elucidating the intrinsic versus task-evoked connection. Here we used computational modeling and graph theoretical metrics to quantify changes in intrinsic functional brain connectivity due to task execution. We used our Large-Scale Neural Modeling framework to embed a computational model of visual short-term memory into an empirically derived connectome. We simulated a neuroimaging study consisting of ten subjects performing passive fixation (PF), passive viewing (PV) and delay match-to-sample (DMS) tasks. We used the simulated BOLD fMRI time-series to calculate functional connectivity (FC) matrices and used those matrices to compute several graph theoretical measures. After determining that the simulated graph theoretical measures were largely consistent with experiments, we were able to quantify the differences between the graph metrics of the PF condition and those of the PV and DMS conditions. Thus, we show that we can use graph theoretical methods applied to simulated brain networks to aid in the quantification of changes in intrinsic brain functional connectivity during task execution. Our results represent a step towards establishing a connection between intrinsic and task-related brain activity.

**Author Summary:** Studies of resting-state conditions are popular in neuroimaging. Participants in resting-state studies are instructed to fixate on a neutral image or to close their eyes. This type of study has advantages over traditional task-based studies, including its ability to allow participation of those with difficulties performing tasks. Further, a resting-state neuroimaging study reveals intrinsic activity of participants’ brains. However, task-related brain activity may change this intrinsic activity, much as a stone thrown in a lake causes ripples on the water’s surface. Can we measure those activity changes? To answer that question, we merged a computational model of visual short-term memory (task regions) with an anatomical model incorporating major connections between brain regions (non-task regions). In a computational model, unlike real data, we know how different regions are connected and which regions are doing the task. First, we simulated neuronal and neuroimaging activity of both task and non-task regions during three conditions: passive fixation (baseline), passive viewing, and visual short-term memory. Then, applying graph theory to the simulated neuroimaging of non-task regions, we computed differences between the baseline and the other conditions. Our results show that we can measure changes in non-task regions due to brain activity changes in task-related regions.

## INTRODUCTION

Recently, there has been significant interest in investigating the relationship between intrinsic and task-evoked brain activity. This interest is driven by the potential to discover information contained in intrinsic brain activity that would reveal the repertoire of functional brain networks used to execute goal-directed tasks (Cole, Bassett, Power, Braver, & Petersen, 2014). Intrinsic and task-evoked activity are strongly interdependent (Bolt, Anderson, & Uddin, 2017) and understanding this interdependence holds the promise of providing a link between resting state and task-based empirical findings (Cole et al., 2014). Furthermore, the establishment of a clear relationship between intrinsic and task brain activity would allow the assessment of task-related brain areas in patients unable to comply with such tasks (Branco et al., 2016; H. Liu et al., 2009)

Neuroimaging studies have shown that performance of a cognitive task alters the intrinsic functional connectivity in non-task related brain regions (Bluhm et al., 2011; Tommasin et al., 2017; Vatansever, Menon, Manktelow, Sahakian, & Stamatakis, 2015). Bluhm and colleagues, for example, found increases in functional connectivity between two “default network” brain regions (posterior cingulate / precuneus and medial prefrontal cortex) and the rest of the brain during a visual working memory task as compared to a passive fixation task. In another study, Tommasin and colleagues found reductions in functional connectivity between brain regions within the “default mode network” (DMN) during an auditory working memory task as compared to an eyes-open resting state (RS) task. Similarly, Vatansever and colleagues found reductions in functional connectivity within DMN brain regions during a motor task as compared to a RS task.

A very powerful tool that has been used to quantify changes in intrinsic functional connectivity due to task execution employs graph theoretical methods (Adams, Shipp, & Friston, 2013; Bolt, Nomi, Rubinov, & Uddin, 2017; Cohen & D’Esposito, 2016; Fuertinger, Horwitz, & Simonyan, 2015; Krienen, Yeo, & Buckner, 2014; Moussa et al., 2011). Graph theoretical metrics have been used in the last decade to study functional and structural brain networks as they provide ways to quantify both global network organization and local network properties (Bolt, Nomi, et al., 2017; Rubinov & Sporns, 2010).

A recent computational study (Lee, Bullmore, & Frangou, 2017) demonstrated the reliability of graph theoretical metrics obtained from simulated intrinsic brain activity. Lee and colleagues modeled brain regions as Kuramoto oscillators coupled by weights extracted from a structural connectome (Hagmann et al., 2008). After finding an optimal functional connectivity matrix (one that resembled the RS empirical connectivity matrix), they set out to compute global and local network metrics and compared them to empirically-obtained graph metrics during the resting state. They found that simulated brain activity can be reasonably used to model graph theoretical metrics of brain organization.

However, there is a need to test the use of graph theoretical metrics on simulated intrinsic activity during task execution. We aimed to use computational modeling and graph theoretical metrics to quantify differences in intrinsic functional brain connectivity of non-task-related brain regions due to increasing task demands. We used a large-scale computational model of visual processing that was previously verified against single-unit recordings in non-human primates and empirical PET, fMRI, and MEG data (Banerjee, Pillai, & Horwitz, 2012; Corbitt, Ulloa, & Horwitz, 2018; Horwitz et al., 2005; Q. Liu, Ulloa, & Horwitz, 2017; Tagamets & Horwitz, 1998; Ulloa & Horwitz, 2016). We embedded the visual processing model in a structural connectome (Hagmann et al., 2008) to examine differences in intrinsic neural activity between three conditions: passive fixation (PF), passive viewing (PV), and a visual delayed match-to-sample (DMS) task. Specifically, we set out to investigate whether computational modeling and graph theoretical metrics could be used to quantify and understand intrinsic neural activity changes in non-task brain regions due to increasing task demands.

## RESULTS

To perform the current study, we embedded a biologically realistic model of visual short-term memory (Tagamets & Horwitz, 1998), shown in Figure 1, into an anatomical skeleton defined by a 998-node structural connectome (Hagmann et al., 2008), shown in Figure 2, using a blend of our large-scale neural model (LSNM) simulator (Ulloa & Horwitz, 2016) and the Virtual Brain (TVB) simulator (Sanz Leon et al., 2013). The visual short-term memory model used here has been previously verified against single-unit recordings in non-human primates (Tagamets & Horwitz, 1998) and empirical PET (Tagamets & Horwitz, 1998), MEG (Banerjee et al., 2012) and fMRI data (Corbitt et al., 2018; Horwitz et al., 2005; Q. Liu et al., 2017). Such a visual model comprises brain regions that are directly involved in performing a delayed match-to-sample (DMS) task for visual objects. As mentioned above, we added a structural connectome to provide neural noise to the simulated neural activity during the DMS task, and in return, to receive inputs back from the DMS task nodes. We have described our framework in a previous paper (Ulloa & Horwitz, 2016) where we focused on the fMRI BOLD signal generation during the DMS task. In the current work, we sought to analyze the functional connectivity (FC) configurations in brain regions not driving task execution. These ‘non-task’ brain regions exhibit intrinsic activity and because of their reciprocal connections with task-specific brain regions, their neural activity can potentially be modulated during task execution.

**Figure 1.**
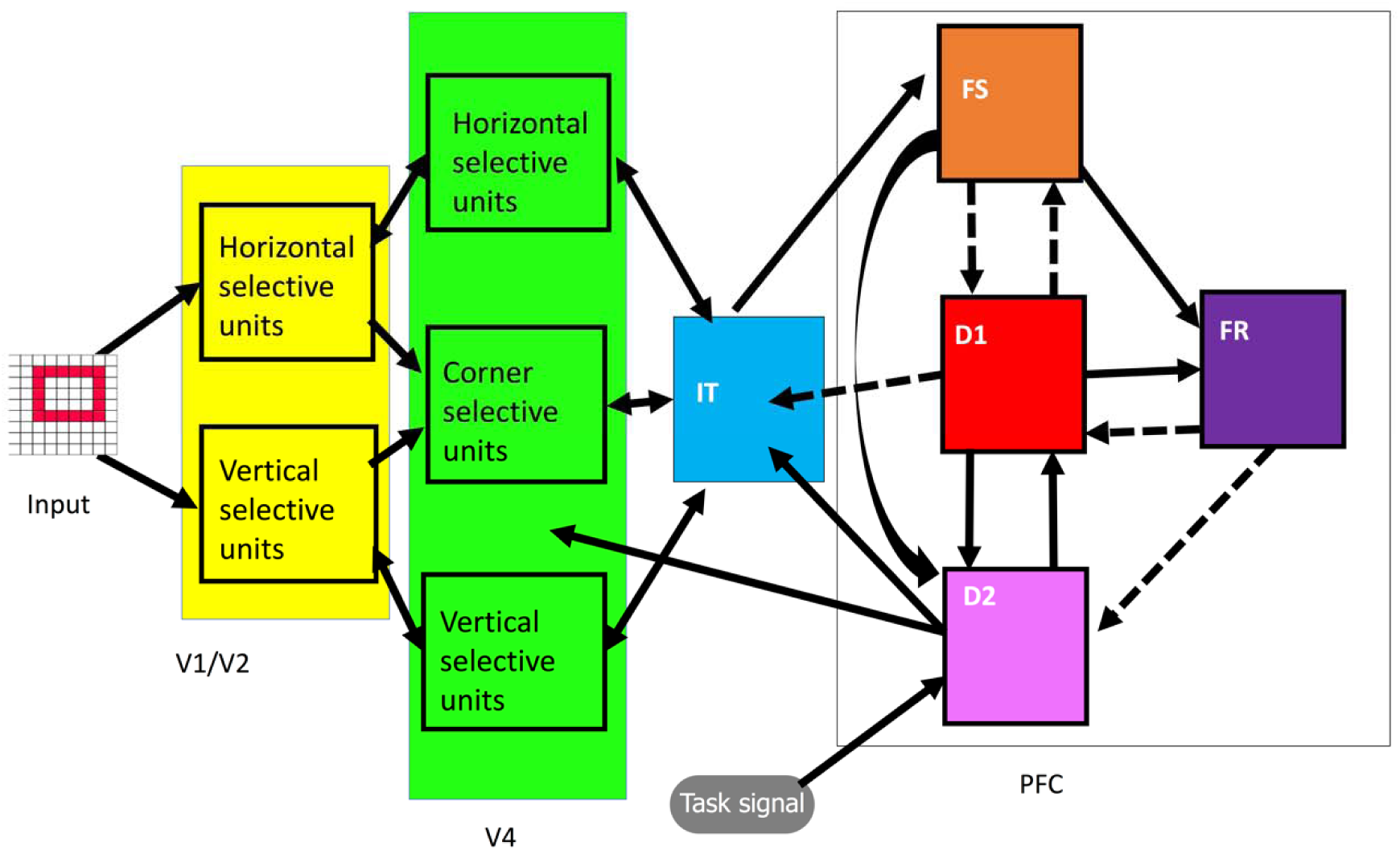
Visual short-term memory model consisted of interconnected neural populations that represent primary and secondary visual (V1/V2, V4), inferotemporal (IT), and prefrontal cortex (PFC). Each one of the sub-modules (shown above as squares) within a given brain module is modeled with 81 (9×9) modified Wilson-Cowan neuronal population units. Solid arrows represent Excitatory to Excitatory connections and dashed arrows represent Excitatory to Inhibitory connections. Adapted from (Horwitz et al., 2005).

**Figure 2.**
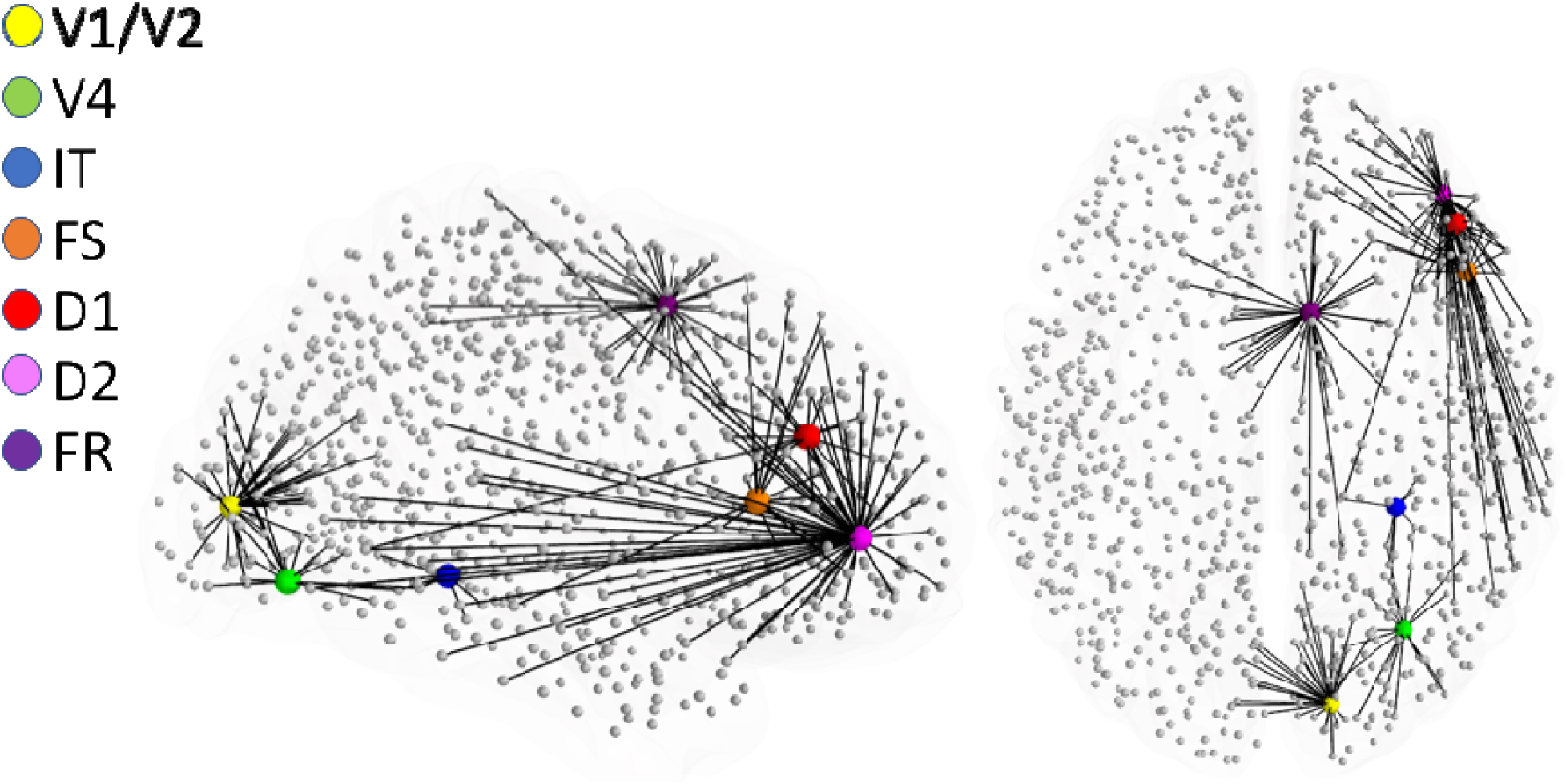
Graphical representation of the location where each of the visual short-term memory nodes was embedded within Hagmann’s connectome (Hagmann et al., 2008). Also shown are direct anatomical connections to connectome nodes from each one of the embedded LSNM nodes.

We generated ten virtual subjects by randomly varying the connection weights among brain regions in the structural visual model (see Methods section for details). We created three experimental conditions: passive fixation (PF), during which simulated subjects with a low “task signal” (roughly equivalent to subjects’ attention level during task execution, but see Methods for definition of this parameter) are fixating on a small dot; passive viewing (PV), during which subjects passively look at visual shapes; and a DMS task, during which subjects compared two shapes presented within 1.5 seconds of each other and responded whether the second shape matched the memory of the first. Each simulated subject performed one 198-second experiment that consisted of 3-trial blocks interspersed with rest blocks (see Methods section for details).

### Changes in simulated BOLD activity of non-task brain regions due to different task conditions

Figure 3 shows typical (averaged across neuronal populations within each brain region) neuronal activity for each condition for task-related brain regions during one trial. Figure 3 shows the task regions increasing activity due to both stimuli presentation (V1, V4, IT, PF), short-term memory maintenance (D1, D2), and response (FR). This increase occurs in the PV and DMS conditions (green and red lines) but not in the PF condition (blue line). Thus, the stimulus used in the PF condition (a small dot) does not generate visible changes in the neuronal activity of task regions. The details of the task-related responses shown in Figure 3 have been discussed in detail in previous papers (Horwitz et al., 2005; Ulloa & Horwitz, 2016). Figure 4 shows the BOLD signal averaged across those brain regions with direct anatomical connections to task regions. Figure 2 shows a graphical depiction of the non-task nodes that are directly connected to task nodes. Notice how BOLD activity increases during the task blocks (shaded areas) and how they do so more prominently during DMS than during PV and during PV than during PF. Also notice how that BOLD activity change is larger for some of the brain regions with direct connections to IT, FS, D1, D2, FR than those regions with direct connections to V1 and V4. This is due to variations in the strength of the connecting weights from task-related nodes to non-task nodes. As we can see in Figure 4, changes in all task-related brain regions correlate with BOLD signal changes in non-task brain regions directly connected to them.

**Figure 3.**
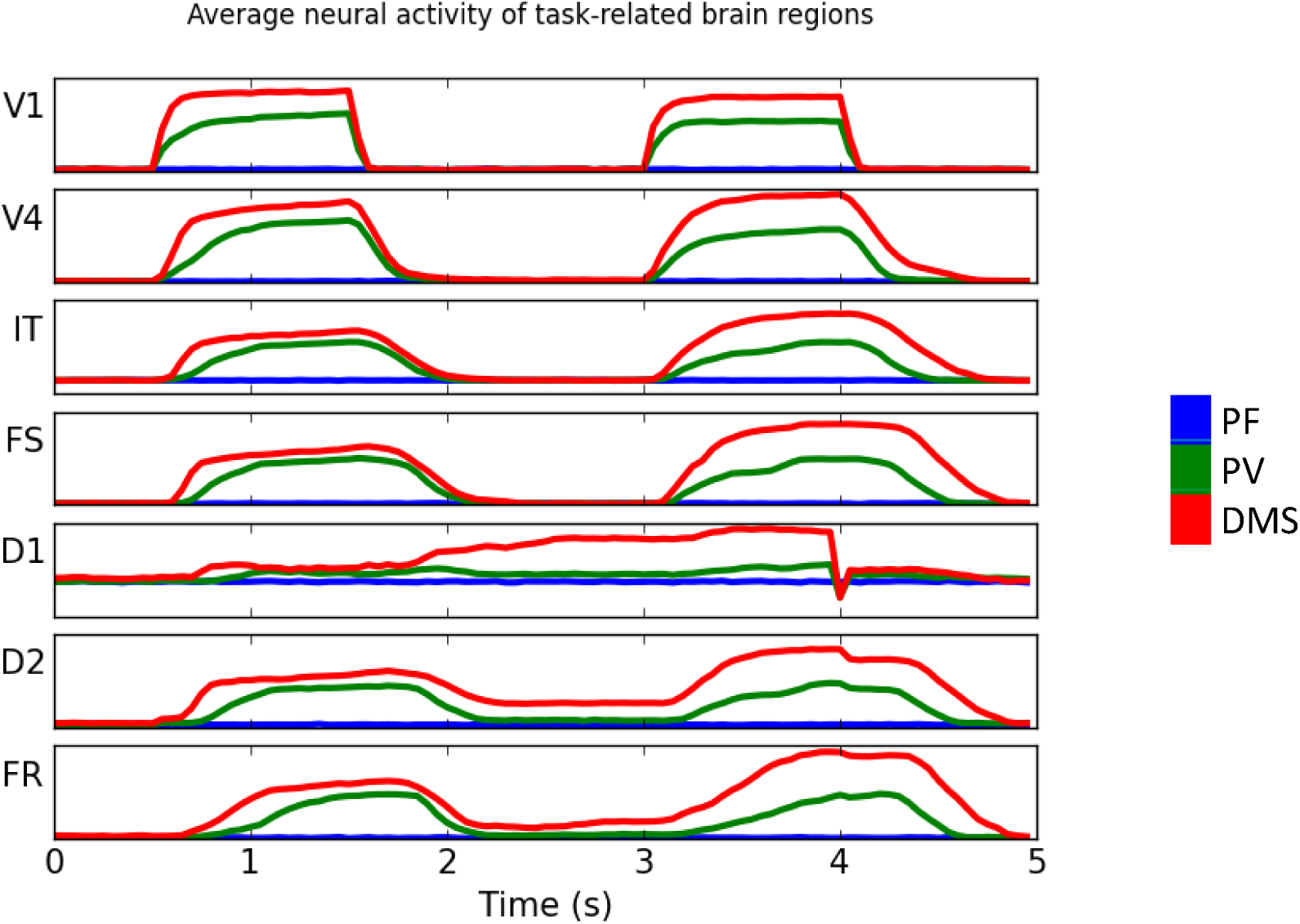
Typical electrical and in neuronal populations of task-related brain regions during one trail of each of the simulated conditions. Key: PF (blue line), PV (green line), DMS (red line). What is shown is the average across all cortical columns in a brain region.

**Figure 4.**
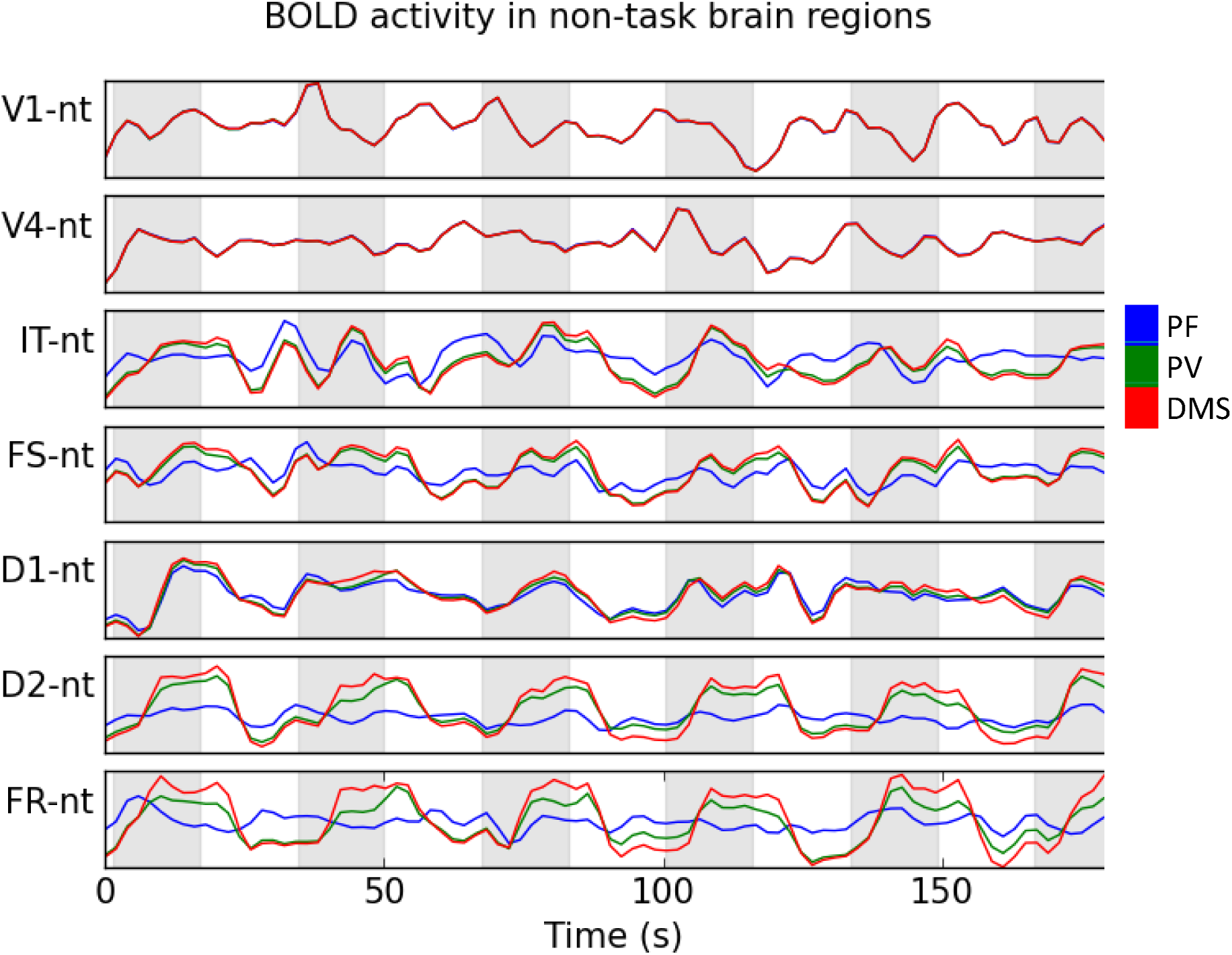
Average BOLD signal of non-task brain regions with direct connections to task related brain regions. A complete trial corresponding to 91 scans is shown above. for the PV and DMS conditions, each experiment above contains 6 task blocks (shaded regions) interspersed with rest blocks.

### Intrinsic FC differences between PF, PV and DMS conditions

We computed FC matrices for the three simulated conditions and for all subjects. Figure 5 shows across-subject averages of FC matrices for the three conditions. Figure 6 shows scatter plots between PF and PV and between PF and DMS conditions. As shown in Figure 6, the correlation coefficients between PF and both PV and DMS were high (0.90 and 0.83, respectively), demonstrating only small differences in the pair-wise consistency of functional connections across conditions. As noted above, these correlation matrices consist only of connectome nodes (e.g., no LSNM task-based nodes were used to construct these matrices). In summary, there were small changes in the pair-wise functional connectivity between PF and PV and between PF and DMS conditions.

**Figure 5.**
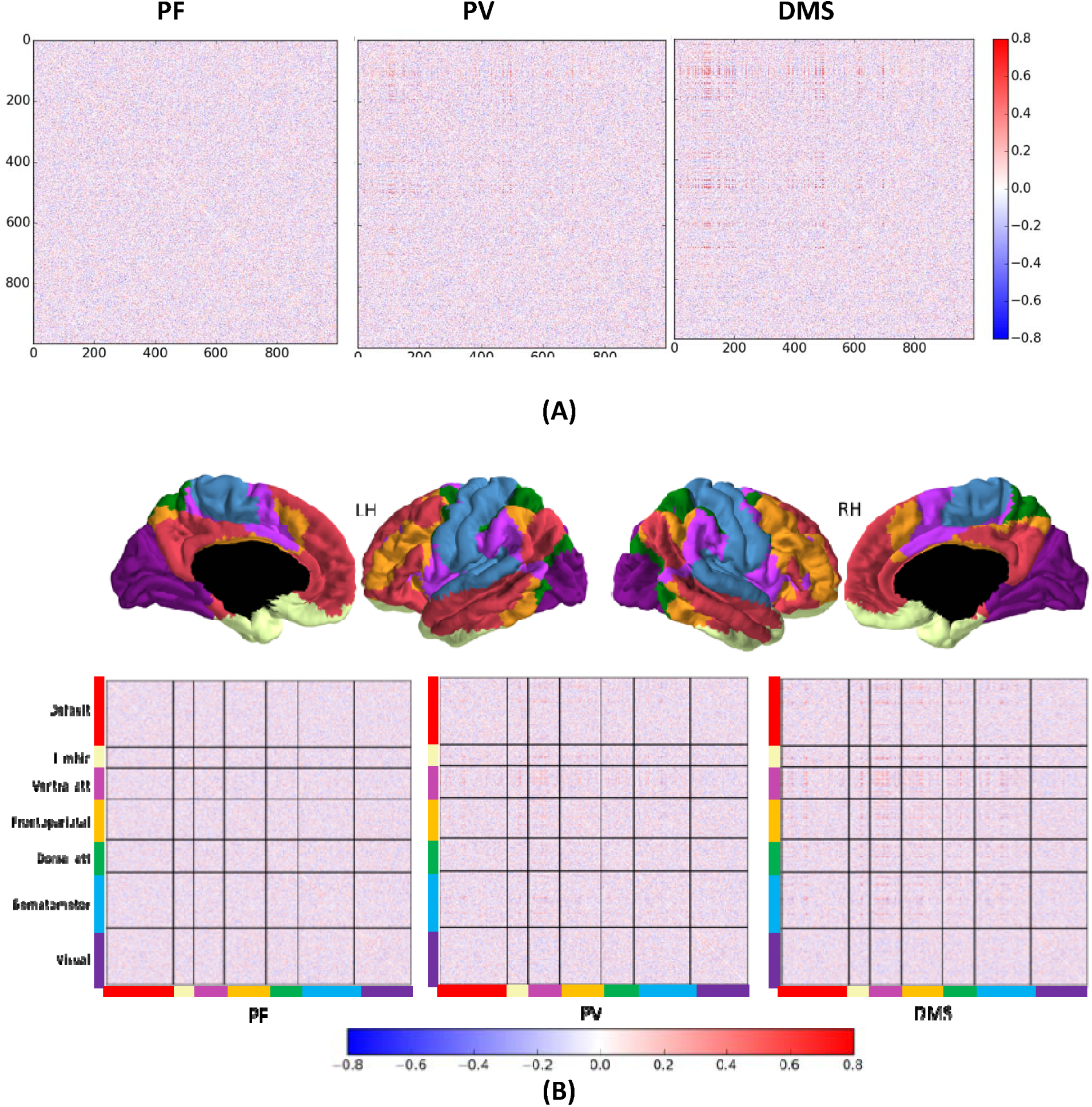
Representative correlation-based functional connectivity matrices for the three conditions simulated. Subject 12 is shown above. (A) The nodes in each matrix are arranged using the standard connectome files in (Hagmann et al., 2008). (B) Nodes in the matrix have been rearranged to match Yeo et al (Yeo et al., 2011) parcellation (7 modules). Brain parcellation was displayed using Freesurfer.

**Figure 6.**
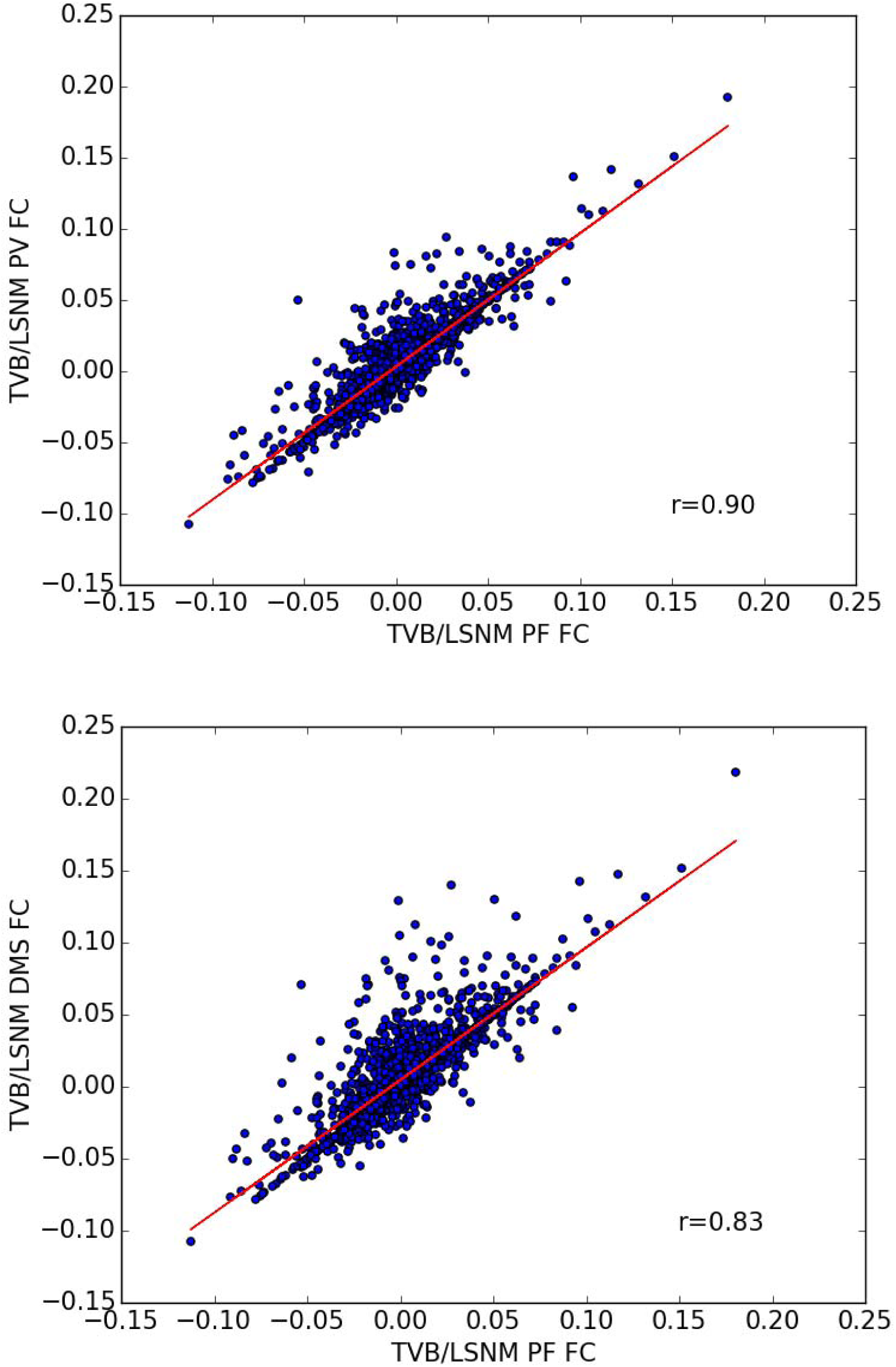
Correlation between PF and PV and between PF and DMS weighted functional connectivity matrices.

### Graph theoretical metrics of PF, PV, and DMS conditions

Using graph theoretical methods (Rubinov & Sporns, 2010), we computed eight network metrics (see Methods section for definition of each metric): global and local efficiencies, average clustering coefficient, characteristic path length, eigenvector centrality, betweenness centrality, participation coefficient, and modularity. We calculated these metrics using weighted FC matrices for a range of plausible threshold densities (Di, Gohel, Kim, & Biswal, 2013). Figure 7 shows across-subject averages of those metrics for a range of network densities (Di et al., 2013). Figure 7 shows that as the task changed from PF to PV to DMS, there was an increase in global efficiency, local efficiency, average clustering coefficient and average betweenness centrality (mostly at the lowest threshold studied, 5%), and modularity. Conversely, as the task changed from PF to PV to DMS, there was a decrease in average characteristic path length, average eigenvector centrality, and average participation coefficient.

**Figure 7.**
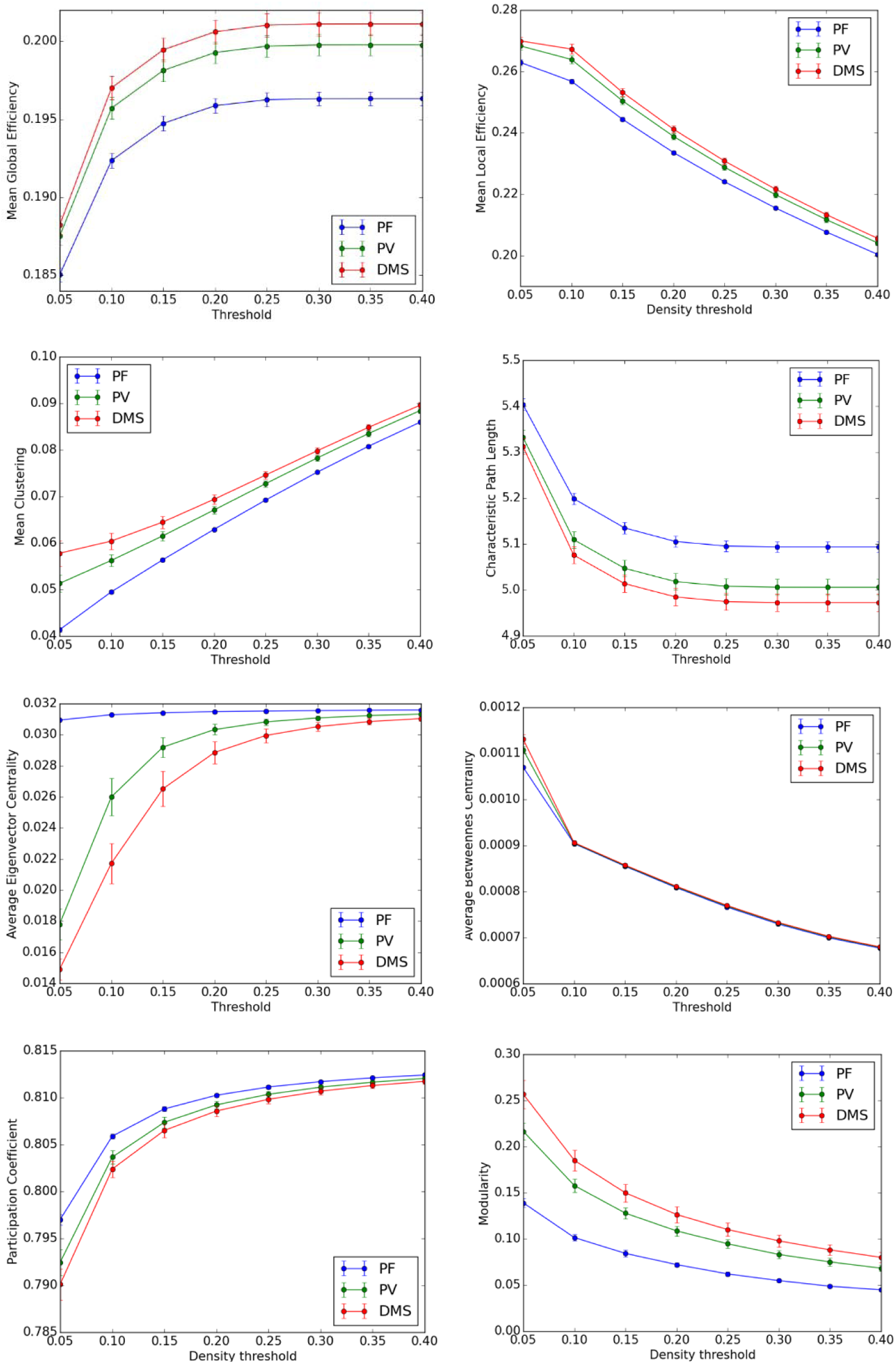
Mean graph theoretical metrics for each condition and for a range of network densities (5 to 40 %). Error bars correspond to standard deviation.

### Differences in graph metrics between PF and PV and between PF and DMS

For each graph metric obtained, we computed the relative difference (see Methods section for details) between PF and PV and between PF and DMS (see Figure 8). We observed significant differences between PF and PV and between PF and DMS in modularity (54.2 ± 8% and 81.3 ± 11.6%, respectively), eigenvector centrality (16.3 ± 1.7% and 22.1 ± 1.8%, respectively) and clustering coefficient (7.9 ± 1.3% and 12.7 ± 2%); smaller changes in global efficiency (1.7 ± 0.2% and 2.4 ± 0.3), local efficiency (2.2 ± 0.3% and 3.2 ± 0.4%), characteristic path length (1.7 ± 0.1% and 2.3 ± 0.3%), betweenness centrality (1.6 ± 0.3% and 2.6 ± 0.4%), and participation coefficient (0.2 ± 0.1% and 0.4 ± 0.1%).

**Figure 8.**
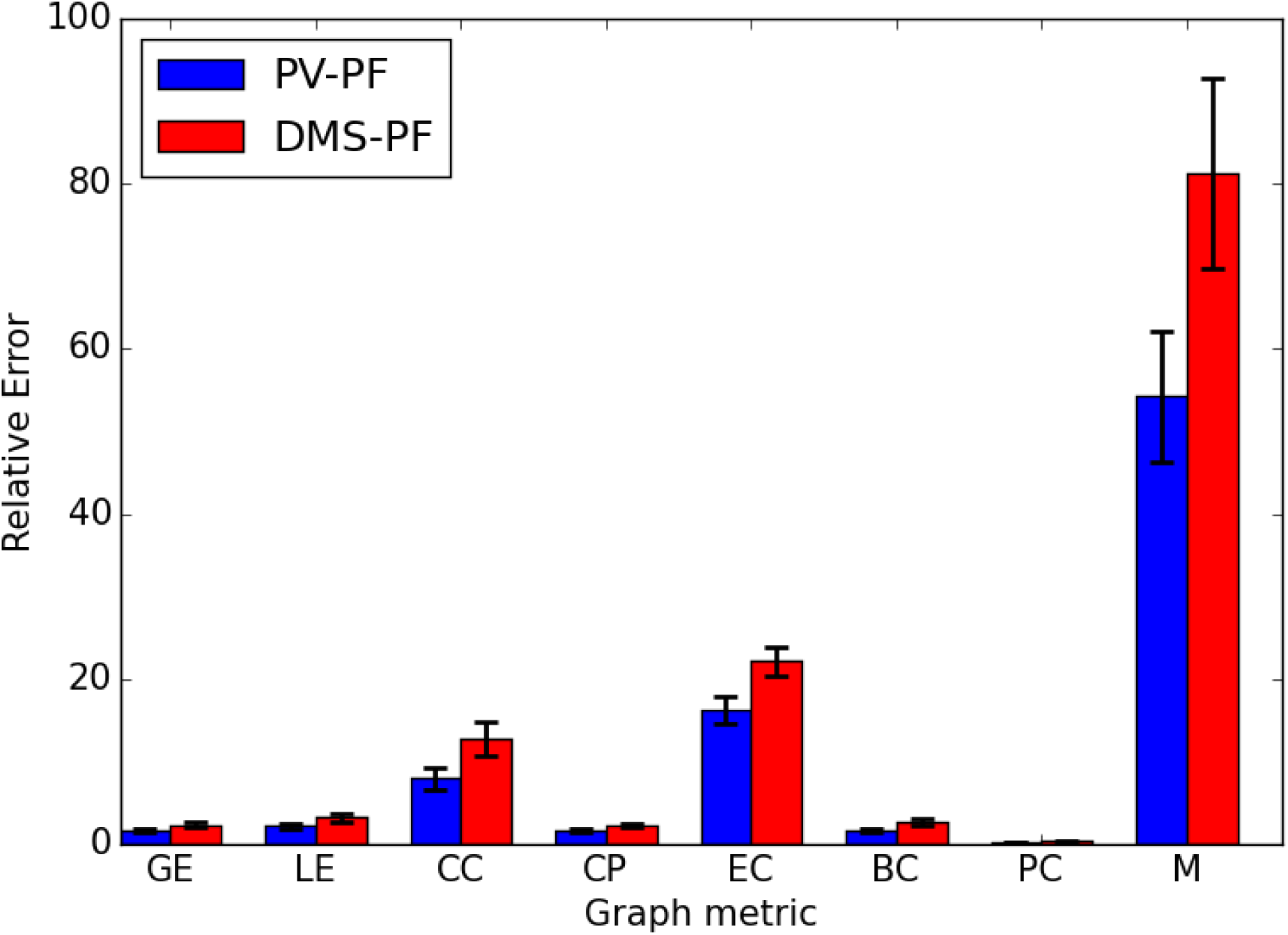
Relative difference between PF and PV and between PF and DMS for each one of the graph metrics in Figure 7. Error bars correspond to standard deviation.

### Differences in modularity between conditions

To further visualize the large differences in modularity configurations during the three simulated conditions, we rendered the binary FC network in each condition as connection space graphs using Gephi (Bastian, Heymann, & Jacomy, 2009); www.gephi.org). We used the algorithm of Blondel et al (Blondel, Guillaume, Lambiotte, & Lefebvre, 2008) to find the modularity at a density threshold of 10%. Figure 10 shows connection space graphs displayed on a radial axis layout (axis have a slight spiral to improve visualization of inter-module connectivity). Nodes that belong to the same module are represented by the same color and group together on the same radial axis. The connections between nodes have the color of the node where those connections originate. We can see a decrease in the number of modules, from 8 in PF to 6 in PV to 3 in DMS and an increase in modularity (see increase in modularity graph in Figure 7). The increase in modularity from PF to PV to DMS means that the functional network rearranges itself into fewer modules with more functional connections between nodes within the same module (compare the very clearly defined modules in DMS versus PF and DMS versus PV in Figure 10). We emphasize again that these results refer to non-task related nodes.

**Figure 10.**
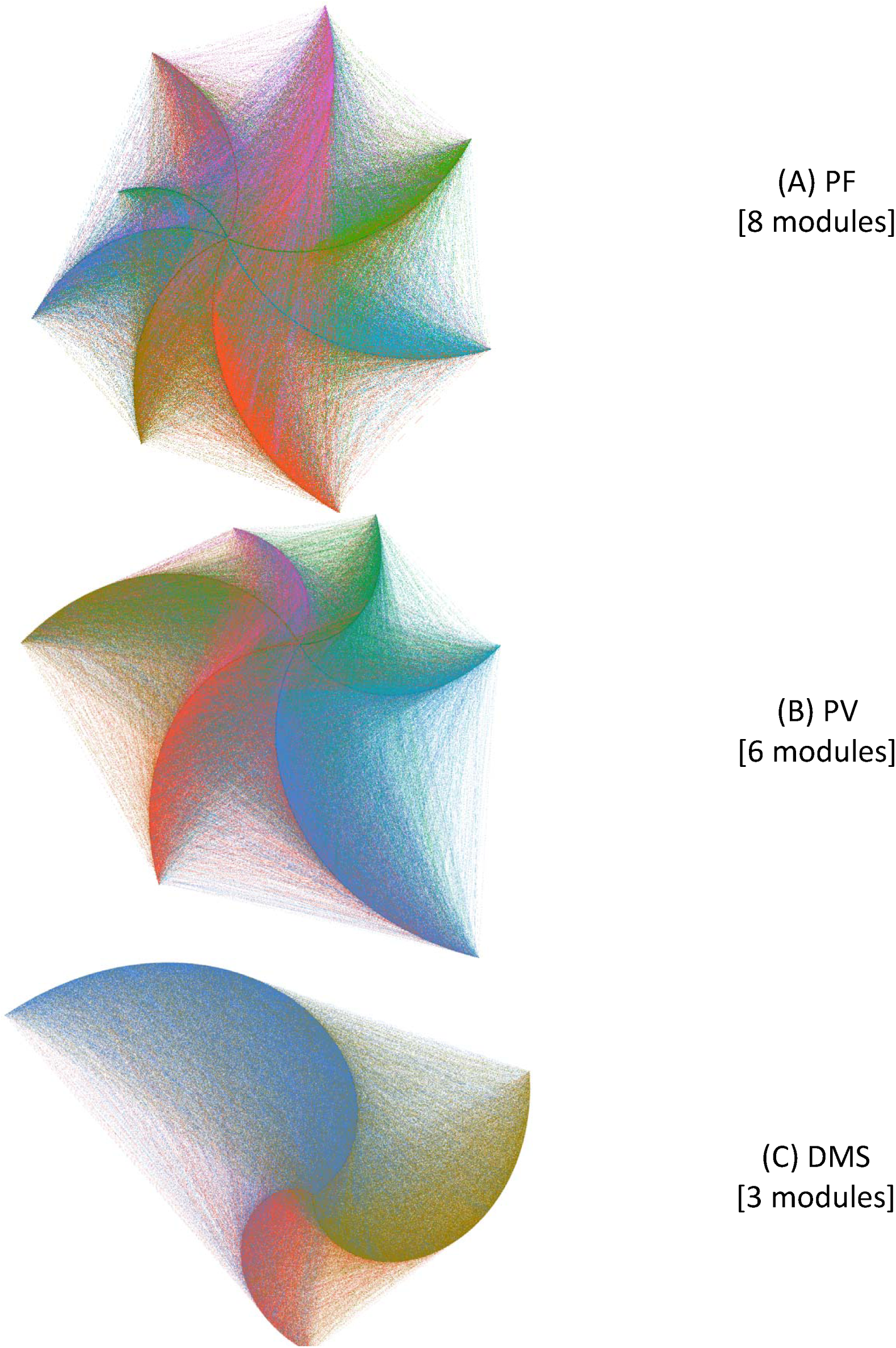
Modular structure of functional connectivity between non-task nodes in conditions (A) PF, (B) PV, and (C) DMS. The graphs used unweighted, undirected functional connectivity matrices at a density threshold of 10 %. These graphs were rendered using the radial axis layout of Gephi (Bastian et al., 2009) and the modular structures were computed using the algorithm of (Blondel et al., 2008).

## DISCUSSION

Using a large-scale computational model of visual short-term memory embedded into an anatomical connectome, we compared simulated intrinsic brain activity of non-task related brain regions during three tasks: passive fixation (PF), during which simulated subjects with a low “task signal” or “attention” level are fixating on visual stimuli (a small dot); passive viewing (PV), during which subjects passively watch changing visual shapes but take no action; and a DMS task, during which subjects compared two shapes presented within 1.5 seconds of each other and responded whether the second shape matched the memory of the first. The PF condition may be considered equivalent to a resting state condition as a passive fixation task has been often used in RS fMRI studies. The key difference between the PF and the PV conditions was that the stimulus during the PF condition was an unchanging small dot whereas in the PV condition several different and larger stimuli were presented. The key difference between the PV and the DMS conditions was the level of the “task” or attention signal, which was set to a low level in the PV condition and to a high level during the DMS condition. As discussed in the Methods section, the task signal level determines whether an input stimulus is going to be retained in short-term memory (Horwitz et al., 2005). Additionally, because of feedback connections from D1 in prefrontal cortex to IT and V4 (see model diagram in Figure 1), the task signal level indirectly influences neuronal activity in V1, V4, and IT (compare neuronal activity in V1, V4, and IT during different conditions in Figure 3).

To quantify differences between PF, PV and DMS conditions, we used pair-wise temporal Pearson correlations (FC matrices) and graph theory metrics of fMRI FC matrices. Whereas we found small differences between the FC matrices of the simulated conditions, these differences we not particularly impressive. However, we found clear-cut differences in each of the graph theory metrics: Graded increases from PF to PV to DMS in global efficiency, local efficiency, clustering coefficient, betweenness centrality and modularity; and graded decreases in the from PF to PV to DMS in characteristic path length, eigenvector centrality, and average participation coefficient. Our simulated graph theory results largely agree with empirical studies, as will be discussed below in detail.

In our computer simulations, the intrinsic brain activity across different conditions is modulated by ongoing neural activity in brain regions engaged in each task (task brain regions). This modulation happens through the strength of the anatomical connections of those brain regions to the rest of the brain (non-task brain regions, see Figure 2).

When the brain engages in a behavioral task, the activity in neuronal populations driving the task has the potential of reverberating throughout the brain, thereby altering the intrinsic neural activity of neuronal populations not involved in the task. A crucial question is whether one can quantify those changes in intrinsic functional connectivity. Computational modeling can be useful in this regard, as it allows us to isolate non-task from task neuronal populations and to convert simulated synaptic activity into neuroimaging time-series which in turn can be converted to FC matrices. Furthermore, unlike empirical data, in a computational model we know which neuronal populations participate in the task and which ones do not.

A commonly used method to simulate the resting state is by modeling local neuronal populations with oscillators and using the structural connections obtained from diffusion tractography as connection weights between the model neuronal populations. A parameter search is then conducted to find a global coupling parameter and a white matter conduction speed producing a simulated FC matrix that best matches an empirical FC matrix (Cabral, Hugues, Sporns, & Deco, 2011; Ghosh, Rho, McIntosh, Kotter, & Jirsa, 2008; Gilson, Moreno-Bote, Ponce-Alvarez, Ritter, & Deco, 2016; Hansen, Battaglia, Spiegler, Deco, & Jirsa, 2015; Honey et al., 2009; Lee et al., 2017; Roy et al., 2014; Sanz-Leon, Knock, Spiegler, & Jirsa, 2015). This is the method we used to generate intrinsic activity in the “rest of the brain” of our simulations.

### Consistency of pair-wise functional connectivity across task conditions

There was a high correlation between the pairs in the FC connectivity matrices between PF and PV and between PF and DMS (Figure 6). Several researchers have used pair-wise spatial correlations between functional connectivity (FC) matrices to compare intrinsic to task-evoked conditions (Bolt, Nomi, et al., 2017; Buckner et al., 2009; Cohen & D’Esposito, 2016; Cole et al., 2014; Di et al., 2013; Krienen et al., 2014; Smith et al., 2009). Generally, there is a relatively high spatial correlation (i.e., 0.64 – 0.9) between a passive condition (such as visual fixation or eyes closed, which are often used to study intrinsic brain activity) and a task condition. Despite such high correlations, differences do exist between passive and task FC, and those differences may be attributable to functional modifications that allow the brain to focus on performing a given task (DeSalvo, Douw, Takaya, Liu, & Stufflebeam, 2014; Di et al., 2013; Tomasi, Wang, Wang, & Volkow, 2014).

Bolt and colleagues (Bolt, Nomi, et al., 2017) recently showed that one can have largely consistent FC between passive and task conditions, and at the same time have largely different whole-brain graph theoretical metrics between passive and task conditions. However, a description of the mechanisms behind those seemingly divergent results has not yet been provided.

### Increases in Global Efficiency

Our study resulted in higher global efficiency for DMS than for PV and for PV than for PF. During the simulated PF condition, the stimuli used is small and mostly activates V1/V2 and V4 and IT areas to a small degree (blue lines in Figure 3), During the PV condition, the larger stimuli used causes an increase of neuronal activity in V1/V2, V4, IT, FS, D1, D2, FR (as shown in the trial time-series of Figure 3, green lines), thereby contributing to an increase in neuronal activity of non-task nodes directly connected to task nodes (see green lines in the shaded areas of the time-series in Figure 4). During the DMS condition, the neuronal activity across the task brain regions is higher than during the PV condition (red lines in Figure 3). This increase in neuronal activity of task brain regions contributes to an increase in neuronal activity of several of the non-task brain regions with direct connections to task regions during PV and DMS conditions as compared to PF condition (see Figure 4). As shown in the FC matrices of Figure 5, there is an increase in the correlation of several pair-wise connections from PF to PV to DMS. This increase in functional connectivity contributed to a consistent increase in global efficiency from PF to PV to DMS (Figure 7).

Graph theoretical measures in empirical studies have consistently shown higher global efficiency during task than during passive conditions (although this could depend on the complexity of the task, but see (Cohen and D’Esposito 2016)). The global efficiency has been found to be higher during a task than during passive fixation (Bolt, Nomi, et al., 2017; Cohen & D’Esposito, 2016), higher during a task than during an eyes closed condition (Fuertinger et al., 2015), greater during a one-back visual memory task than during passive viewing and an eyes closed condition (Wen et al., 2015), and higher for coactivation studies than during RS (Di et al., 2013). In our simulations, the global efficiency is higher during DMS than during PV and PF. This is due to the short-memory task causing an increase of neural activity in brain regions that are in turn connected to a widely distributed network in the rest of the brain.

### Increases in Local efficiency

Our simulations showed a greater local efficiency for DMS than for PV and for DMS than for PF. This is consistent with empirical studies showing an increase in local efficiency with increasing task demands (Wen et al., 2015).

### Increases in Clustering Coefficient

Our simulations showed a greater clustering coefficient during DMS than during PV and during PV than during PF. Previous empirical studies have found a clustering coefficient that is greater for task than during passive fixation (Bolt, Nomi, et al., 2017), lower during a blend of activation studies than during resting state (Di et al., 2013), and greater during a language task than during eyes closed (Fuertinger et al., 2015).

### Increases in characteristic path length

Our simulations showed smaller characteristic path length during DMS than during PV and during PV than during PF. This is to be expected because as the global efficiency increases, the characteristic path length decreases.

### Decreases in mean Eigenvector Centrality

Our simulations showed smaller eigenvector centrality during DMS than during PV and during PV than during PF. The eigenvector centrality metric provides a measure of how well-connected a given node is considering how well connected that node’s neighbors are. Thus, eigenvector centrality is recursive because a given node’s eigenvector centrality depends on the node’s neighbors’ eigenvector centrality. To get a more detailed view of the reason behind smaller mean eigenvector centrality for more complex tasks (Figure 7), we rendered the eigenvector centrality for each node on axial and sagittal views of the brain (Figure 9A). Figure 9A shows that as the task complexity increases (from PF to PV to DMS) the eigenvector centrality increases in a few nodes and decreases in most other nodes. Thus, on average the eigenvector centrality decreases but the nodal eigenvector centrality in a few nodes increases as the task complexity increases. Note that several of the nodes in which the eigenvector centrality increases during PF and DMS are the nodes that are directly connected to task nodes (compare to Figure 2). The reason the increases are concentrated on the right side of the brain is due to the task nodes, which are embedded in the right side of the brain, having direct connections mostly to the right side of the brain (see Figure 2). Compare the changes in eigenvector centrality with the changes in betweenness centrality (Figure 7) which remain almost the same during PF, PV and DMS (Figure 9B).

**Figure 9.**
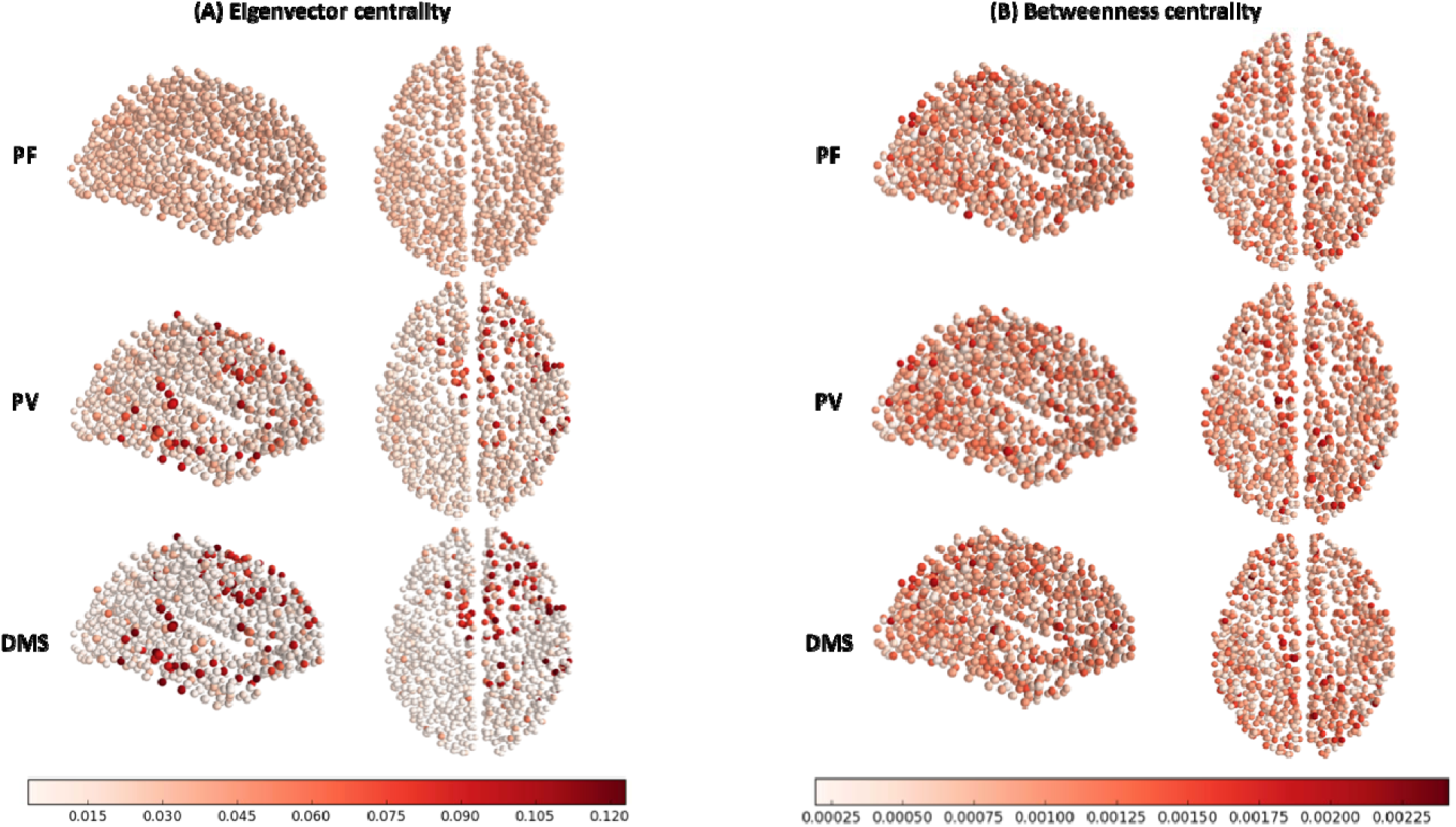
Eigenvector centrality (A) and betweenness centrality (B) depicted on a node-by-node basis on sagittal (left) and axial (right) views of the brain. The density threshold used for the depiction above was 10 %.

### Increases in Betweenness Centrality

Our simulations show a higher betweenness centrality at the lower density threshold (5%) but the average betweenness centrality is very similar across all the other density thresholds (Figure 7). As mentioned above, the betweenness centrality at each individual node (Figure 9B) remains relatively constant across conditions. Previous empirical studies have shown a difference in nodal centrality when resting state and task are compared (Di et al., 2013).

### Decreases in Participation Coefficient

Our simulations showed greater participation coefficient (in a predefined set of modules) for PF than for PV and for PV than for DMS (Figure 7). Participation coefficient measures each node participation in a set of predefined modules. We used the modules defined by Hagmann et al (Hagmann et al., 2008). Previous studies have shown a higher participation coefficient (between-module connectivity) during passive fixation than during a semantic task (DeSalvo et al., 2014).

### Increases in Modularity

Our simulations showed a smaller modularity for PF than for PV and for PF than for DMS. Some empirical studies have found a greater modularity metric during RS than during a blend of activation studies (Di et al., 2013), and a greater modularity during passive fixation than during an n-back task using visually-presented phonemes (Cohen & D’Esposito, 2016). However, Cohen et al (Cohen & D’Esposito, 2016) found a similar modularity during passive fixation and a finger tapping task. Other empirical studies have found that that the modularity varies as a function of performance, but here the evidence is also inconsistent. For example, Stevens et al (Stevens, Tappon, Garg, & Fair, 2012) found a positive correlation between RS modularity and visual working memory capacity and Meunier et al (Meunier et al., 2014) found a negative correlation between modularity and memory scores in an odor recognition task. Additionally, Yue et al (Yue et al., 2017) have found significant individual variability in modularity during resting state.

### Related computational studies comparing resting state and task-based functional connectivity

Two previous computational approaches have compared the intrinsic brain activity obtained during resting state versus the one obtained during task; however, none of those models was specifically concerned with quantifying intrinsic activity differences between different task conditions (which is the goal of our paper). The first one of those studies, by Ponce-Alvarez and colleagues (Ponce-Alvarez, He, Hagmann, & Deco, 2015) simulated RS using a set of mean field equations (excitatory-inhibitory pairs) interconnected by the anatomical connections of a 66-node connectome. A visual task was approximated by applying external stimulation (stationary inputs) to visual nodes during the RS simulation. Ponce-Alvarez’s model revealed a decreased synaptic activity variability during the visual task as compared to the RS condition.

The second computational study comparing task versus rest (Cole, Ito, Bassett, & Schultz, 2016) similarly applied stationary inputs to a set of neighboring nodes in a simplified computational model to simulate six different tasks. Cole and colleagues used the FC strengths during a passive task to predict the fMRI task activation of a held-out brain region. They did this for each one of the brain areas simulated to produce a prediction of the fMRI activity in each one of the brain areas simulated given a passive task FC matrix.

### Caveats and limitations of our study

Different passive experimental conditions have been used in neuroimaging to study intrinsic brain activity (also referred to as the “resting state (RS)”) (Biswal, Yetkin, Haughton, & Hyde, 1995; Fox, Corbetta, Snyder, Vincent, & Raichle, 2006; Greicius, Krasnow, Reiss, & Menon, 2003). Three of the conditions most commonly used as a resting state condition are passive fixation (PF), eyes open with no fixation, and eyes closed. Yan and colleagues (Yan et al., 2009) found significantly higher FC in Default Mode Network (DMN) brain areas during eyes open than during eyes closed condition. It is also important to emphasize that the functional magnetic resonance (fMRI) results can vary depending on several other factors including: how a RS task is defined (Van Dijk et al., 2010; Yan et al., 2009), which task instructions are given to subjects (Benjamin et al., 2010), and whether subjects were engaged in a task prior to RS (Waites, Stanislavsky, Abbott, & Jackson, 2005). Thus, whereas one can compare (within the limitations outlined below) the results of our study with empirical studies using passive fixation, our results cannot be directly extrapolated to all RS-fMRI studies.

One way in which the simulations presented here are different from our previous paper (Ulloa & Horwitz, 2016) is that the model response units have been relocated from prefrontal cortex to PreSMA. The relocation of the response units to PreSMA is based on an fMRI study by (Pessoa, Gutierrez, Bandettini, & Ungerleider, 2002), who found an increase in BOLD fMRI in the PreSMA area at the end of the delay period during a visual working memory task. Additionally, a study by (Petit, Courtney, Ungerleider, & Haxby, 1998) has also demonstrated BOLD fMRI activity in the PreSMA area during a working memory task. The relocation from previous studies from our lab of the model response units to PreSMA makes biological sense as it better reflects the complexity of the task we are trying to simulate. The identification of realistic locations within the brain for each one of the model units is crucial as different locations of task-related modules will modulate different non-task nodes in the connectome, thereby producing different FC configurations.

One of the limitations of our study is that our model connectome does not have other sensory systems apart from the visual system. Therefore, one should exercise caution when comparing FC matrices of our simulation to empirical ones as the empirical ones would contain higher FC that are the result of other sensory systems being activated by either intrinsic or extrinsic processes. For example, in an fMRI scanner room, there is significant auditory stimulation (scanner noise) as well as somatosensory input, which we have not simulated in the present work.

In our simulations, we only embedded the visual model in the right hemisphere. As a result, the intrinsic activity was mostly localized to the right hemisphere. Nonetheless, there were significant intrinsic activity changes in the left hemisphere, and those were caused by structural connectivity between both hemispheres.

Another limitation of our study is that the weights of the structural connectome used in this paper are undirected and we assumed all connection weights to be excitatory. It is well known that diffusion tractography has serious limitations as it produces a significant number of false positives (Maier-Hein et al., 2017), has relatively low resolution and measures white tracts only indirectly (Jbabdi, Sotiropoulos, Haber, Van Essen, & Behrens, 2015). Some researchers have simulated whole brain activity using connectome datasets obtained from reconstructions of retrograde tracer injections in macaques (Chaudhuri, Knoblauch, Gariel, Kennedy, & Wang, 2015) or a composite of diffusion spectrum imaging in humans and macaque tracer data (Sanz-Leon et al., 2015). Despite the low resolution and lack of sign and direction of the human tractography data, we decided to use it as it allowed the “brain regions” of our task-based simulator to be embedded into plausible locations within the structural connectome.

## CONCLUSIONS

In conclusion, we used our large-scale neural modeling framework to quantitatively compare neural dynamics of non-task brain regions during passive fixation, passive viewing, and a visual short-term memory task. We were able to obtain quantitative measures of differences in simulated functional connectivity by using graph theoretical methods. Our simulated graph theory results largely agreed with experiments. We were also able to relate those network-level changes to the underlying model mechanisms. We showed that we can use computational modeling, functional connectivity and graph theoretical metrics to quantify changes in intrinsic FC of non-task brain regions due to increasing task demands. Our work is relevant to the characterization of intrinsic brain activity differences between passive and active task conditions and to the use of neural modeling in the design of empirical studies and the comparison of competing hypothesis of brain function.

## METHODS

In the present work, we analyzed functional connectivity derived from BOLD fMRI time-series, calculated from simulated neural activity data using the framework presented in a previous paper (Ulloa & Horwitz, 2016). Whereas in our previous paper we evaluated the FC between brain regions directly involved in executing a task, in the present paper we examined the intrinsic FC in the rest of the brain (brain regions not involved in task execution). To better address that question, we performed a model parameter search to find a reasonable match between empirical and model FC. Below we briefly describe the components of the framework and how it was used to generate the simulated multi-subject experiment presented in this study. The source code of our modeling work, including simulation, analysis and visualization scripts, is freely available at https://nidcd.github.io/lsnm_in_python/.

### Visual object processing model and The Virtual Brain

#### a. Visual object processing model

Our in-house visual (Tagamets & Horwitz, 1998) object processing model consists of interconnected neuronal populations representing the cortical ventral pathway that has been shown to process primarily the features of a visual object. This stream begins in striate visual cortex, extends into the inferior temporal lobe and projects into ventrolateral prefrontal cortex (Haxby et al., 1991; McIntosh et al., 1994; Ungerleider & Mishkin, 1982). The regions that comprise the visual model include ones representing primary and secondary visual cortex (V1/V2), area V4, anterior inferotemporal cortex (IT), and prefrontal cortex (PFC) (see Fig. 1). Each of these regions contain one or more neural populations with different functional attributes (see caption to Fig. 1 for details). This model was designed to perform a short-term memory delayed match-to-sample (DMS) task during each trial of which a stimulus S1 is presented for a certain amount of time, followed by a delay period in which S1 must be kept in short-term memory. When a second stimulus (S2) is presented, the model must respond as to whether S2 matches S1. The model can also perform control tasks: passive fixation (PF) and passive perception of the stimuli (PV), in which no response is required. Multiple trials of the active and passive tasks constitute a simulated functional neuroimaging study.

The key feature used to define a visual object was shape. Model neurons in V1/V2 and V4 were assumed to be orientation selective (for simplicity, horizontal and vertical orientations were used). The structural submodels employed were based on known monkey neuroanatomical data. An important assumption for the visual model, inferred from such experimental data, was that the spatial receptive field on neurons increased along the ventral processing pathway (see (Tagamets & Horwitz, 1998) for details).

Each neuronal population consisted of 81 microcircuits, each representing a cortical column. The model employed modified Wilson-Cowan units (an interacting excitatory and inhibitory pair of elements for which spike rate was the measure of output neural activity) as the microcircuit (Wilson & Cowan, 1972). The input synaptic activity to each neuronal unit can also be evaluated and combinations of this input activity were related to the fMRI BOLD signals via a forward model.

In an earlier version of the model (Horwitz et al., 2005), half the neural populations within the model were ‘non task-specific’ neurons that served as noise generators to ‘task500 specific’ neurons that processed shapes during the DMS task. The model generated time series of simulated electrical neuronal and synaptic activity for each module that represents a brain region. The time series of synaptic activity, convolved with a hemodynamic response function, was then used to compute simulated fMRI BOLD signal for each module representing a brain region, as well as functional connectivity among key brain regions (see (Horwitz et al., 2005) for details on this method). This model was able to perform the DMS task, generate simulated neural activities in the various brain regions that matches empirical data from non-human preparations, and produces simulated functional neuroimaging data that generally agree with human experimental findings (see (Tagamets & Horwitz, 1998) and (Horwitz et al., 2005) for details). In the current paper, we employ the version of the model introduced by Ulloa and Horwitz (Ulloa & Horwitz, 2016) in which non task-specific neurons are replaced by noise-generated activity from neural elements in The Virtual Brain software simulator (Sanz Leon et al., 2013).

#### b. The Virtual Brain

The Virtual Brain (TVB) software (Sanz Leon et al., 2013; Sanz-Leon et al., 2015) is a simulator of primarily resting state brain activity that combines: (i) white matter structural connections among brain regions to simulate long-range connections, and (ii) a given neuronal population model to simulate local brain activity. It also employs forward models that convert simulated neural activity into simulated functional neuroimaging data. TVB source code and documentation are freely available from https://github.com/the-virtual-brain.

In the current paper, for the structural model, we chose the DSI-based connectome described by (Hagmann et al., 2008), which contains 998 nodes. For the neural model for each node, we employed Wilson-Cowan population neuronal units (Wilson & Cowan, 1972) to model the local brain activity because our in-house LSNM simulators use modified Wilson-Cowan equations as their basic neuronal unit. Our forward model that converts simulated neural activity into simulated fMRI is a modification of the Balloon-Windkessel model of Friston et al. (Friston, Mechelli, Turner, & Price, 2000; Stephan, Marshall, Penny, Friston, & Fink, 2007) that is included in the TVB.

### Integrating TVB and LSNM

To perform our computational study, we concurrently ran two neural simulators: Our Large-Scale Neural Model (LSNM) simulator, which generated task-driven neural activity of the brain regions directly involved in the visual DMS task, and The Virtual Brain simulator (TVB) (Sanz Leon et al., 2013) to generate resting-state neural activity in the brain regions not involved in the task. Because the task-based brain nodes were embedded within resting-state brain ROIs, we expected that the neuroimaging activity in key connectome ROIs would differ between passive fixation (PF), passive viewing (PV), and task-based simulations. Here, we sought to quantify those differences, first by comparing the pattern of functional connectivity across conditions, then by using graph theoretical methods to quantify those differences.

Within the LSNM, connections and parameter choices closely follow those in the original papers. Likewise, the connections and parameter choices among TVB nodes closely follow those described by Sanz-Leon et al. (Sanz-Leon et al., 2015). There are two differences between the simulations presented in this paper and the previous (Ulloa & Horwitz, 2016) paper: The location of the FR units has been changed to PreSMA and the global coupling parameter has been changed (after a parameter search procedure detailed below).

#### a. Task-based model node placement in the TVB

The connectome derived by Hagmann and colleagues (Hagmann et al., 2008) serves as a source of neural noise to our task-based neural model. Such a connectome was obtained by averaging the weighted network of five experimental subjects, where each one of the 998 nodes represents a region of interest covering a surface area of approximately 1.5 cm^2^. The connection weights among the nodes represent cortico-cortical connections given by white matter connection density among the given nodes. As stated above, each node is represented by a Wilson-Cowan population unit and thus each node is assumed to be comprised of one excitatory and one inhibitory neural population. We implemented noise as an additive term to the stochastic Euler integration scheme provided by the TVB software.

The locations of the four PFC nodes (FS, D1, D2, FR) require some comment. The inclusion of these four neural populations in the original LSNMs was based on the electrophysiological studies of Funahashi et al. (Funahashi, Bruce, & Goldman-Rakic, 1990) that found in monkey PFC four distinct neuronal responses during a delayed response task: neurons that (1) increased their activity when a stimulus was present (FS), (2) increased their activity during the delay part of the task (D1), (3) increased their activity during both when a stimulus was present and during the delay period (D2), and (4) increased their activity prior to making a correct response (FR). It is not known if these neuronal types are found in separate anatomical locations in PFC or are intermixed within the same brain area, although the latter is the more likely case (except possibly for the FR population). In the original modeling studies of Tagamets and Horwitz (Tagamets & Horwitz, 1998) and Husain et al. (Husain, Tagamets, Fromm, Braun, & Horwitz, 2004), the functional neuroimaging data represented a single region that included all four nodes. To illustrate the integrated synaptic activity and fMRI signal for each one of the modules of the combined LSNM / TVB model separately, we have assigned a different spatial location to each one of the four PFC sub-modules. We have used the Talairach coordinates of the prefrontal cortex, based on (Haxby et al., 1991), for the submodule D1 and have designated spatial locations in adjacent regions of interest for the FS and D2 submodules. The FR submodule has been allocated to a spatial location determined by an fMRI study of working memory in humans (Pessoa et al., 2002). See Table 1 for coordinate locations of each module/submodule of the visual short-term memory nodes within the structural connectome.

**Table 1.** Hypothesized locations, in Talairach coordinates, s, of visual LSNM modules, along with the closest node in the Hagmann et al. connectome. Note that the locations of FS and D2 are not explicitly known (see text) and were chosen only to demonstrate validity of the method.

| Visual submodule | Talairach location | Source | Host connectome node |
| --- | --- | --- | --- |
| V1/V2 | (18, -88, 8) | (Haxby, Ungerleider, Horwitz, Rapoport, & Grady, 1995) | (14, -86, 7) |
| V4 | (30, -72, -12) | (Haxby et al., 1995) | (33, -70, -7) |
| IT | (28, -36, -8) | (Haxby et al., 1995) | (31, -39, -6) |
| FS | Location selected for illustrative purposes |  | (47, 19, 9) |
| D1 | (42, 26, 20) | (Haxby et al., 1995) | (43, 29, 21) |
| D2 | Location selected for illustrative purposes |  | (42, 39, 2) |
| FR | (1, 7, 48) | (Pessoa et al., 2002) | (8, 6, 50) |

#### b. Simulating electrical activity and fMRI activity

##### Electrical activities of each node in Hagmann’s connectome (TVB equations)

Each one of the nodes in Hagmann’s connectome is represented as a Wilson-Cowan model of excitatory (E) and inhibitory (I) neuronal populations, as described in Sanz-Leon et al. (Sanz-Leon et al., 2015):

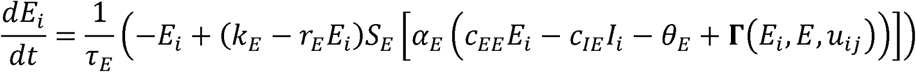

and

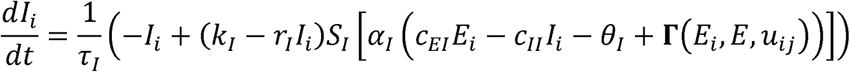

where *S_E_* and *S_I_* are sigmoid functions described by

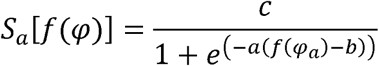

*C_EE_, C_EI_, C_II_, C_IE_* are the connections within the single neuronal unit itself; note that, although the original TVB Wilson-Cowan population model allows us to consider the influence of a local neighborhood of neuronal populations, we have not used this feature in our current simulations and have left that term out of the equations above; **Γ**(*E_K_, E, u_kj_*) is the long-range coupling function, defined as

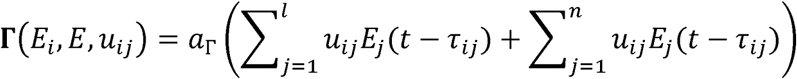

where *l* is the number of nodes in the connectome and *n* is the number of LSNM units connected to a connectome node; *a*_Γ_ is a global coupling parameter (see Supplementary Table S1 and Table S2 for the definition and value of the parameters in the above equations).

##### Electrical activities of each LSNM unit

Each one of the submodules of the LSNM model contains 81 neuronal population units. Each one of those units is modeled as a Wilson-Cowan population of excitatory (*E*) and inhibitory (*I*) elements. The electrical activities of each one of those elements at time *t* is given by the following equations:

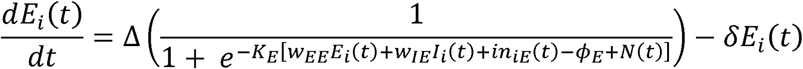

and

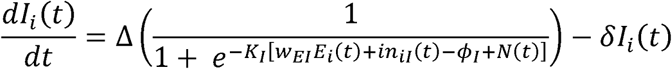

where Δ is the rate of change, *δ* is the rate of decay, *K_E_*, *K_I_* are gain constants, *ϕ_E_*, *ϕ_I_* are input threshold values, *N*(*t*), is a noise term, *w_EE_*, *w_IE_*, *w_EI_* are the weights within a unit (the values of Δ, *δ, K,τ,N* are given in the Supplementary Table S3); *in_iE_*(*t*), *in_iI_*(*t*) are the inputs coming from other brain regions at time *t*. *in_iE_*(*t*) is given by:

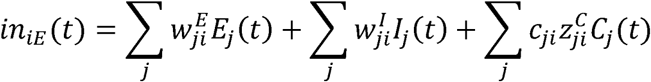

where 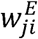 and 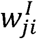 are the weights originating from excitatory (E) or inhibitory (I) unit j from another LSNM unit into the *ith* excitatory element, *C_j_* is the connectome excitatory unit *j* with connections to the LSNM unit *i*, and 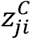 is the value of the anatomical connection weight from connectome unit *j* to LSNM unit *i*, and c_ij_ is a coupling term, which was obtained by using Python’s Gaussian pseudo-random number generator (random.gauss), using *a*_Γ_/81 as the mean value. The input coming into the *ith* inhibitory element, *in_iI_*(*t*), is given by:

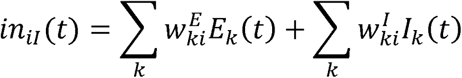

where 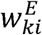 and 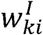 are the weights originating from excitatory (E) or inhibitory (I) unit *k* from another LSNM unit into the ith inhibitory element. Note that there are no connections from the connectome to LSNM inhibitory units. See Supplementary Tables S4 and S5 for details. Note also that, whereas TVB simulator incorporates transmission delay among the connectome nodes, the LSNM nodes do not.

##### Integrated synaptic activity

Prior to computing fMRI BOLD activities we compute the synaptic activity, spatially integrated over each LSNM module (or connectome node) and temporally integrated over 50 milliseconds as described by (Horwitz & Tagamets, 1999)

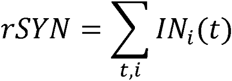

where *IN_i_*(*t*) is the sum of absolute values of all inputs to both *E* and *I* elements of unit *i*, at time *t*, and is given by:

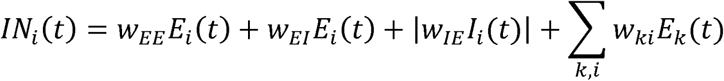

Note that the first three terms above are the synaptic weights from within unit i and the last term is the sum of synaptic connections originating in all other LSNM units and connectome nodes connected to unit i. Note also that, in our current scheme, there are no long-range connections from inhibitory populations.

##### Generation of subjects and task performance of the LSNM model

We generated simulated subjects by creating several different sets of connection weights among submodules of the LSNM visual network until we obtained the number of desired subjects whose task performance was above 60 percent. However, the weights among the nodes with the TVB connectome remained unchanged across subjects. The generation of different connectome sets to simulate individual subjects is outside the scope of the current paper but will be essential for future simulation studies investigating the effects of a behavioral task on non-task brain nodes. Task performance was measured as the proportion of correct responses over an experiment. A response in the response module (FR, described in the caption to Fig. 1) was considered a correct response in each trial if at least 2 units had neuronal electrical responses above a threshold of 0.7 during the response period. To create different sets of weights that were different from the ideal subject, we multiplied feedforward connections among modules in the LSNM visual model by a random proportion of between 0.95 and 1.

##### Equations for the forward fMRI BOLD model

We implemented the BOLD signal model described by (Stephan et al., 2007). We use the output of the integrated synaptic activity above as the neural state equation to the hemodynamic state equations below. The BOLD signal for each region of interest, *y(t*), is computed as follows:

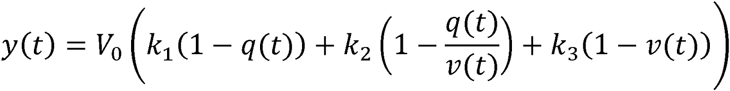

where the coefficients *k*_1_, *k*_2_, *k*_3_ are computed as:

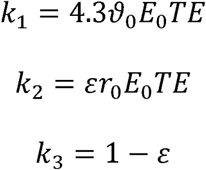

where *V*_0_ is the resting venous blood volume fraction, *q* is the deoxyhemoglobin content, *v* is the venous blood volume, *E_0_* is the oxygen extraction fraction at rest, *ε* is the ratio of intra- and extravascular signals, and *r*_0_-is the slope of the relation between the intravascular relaxation rate and oxygen saturation, *ϑ*_0_ is the frequency offset at the outer surface of the magnetized vessel for fully deoxygenated blood at 3T, and TE is the echo time. The evolution of the venous blood volume *v* and deoxyhemoglobin content *q* is given by the balloon model hemodynamic state equations, as follows:

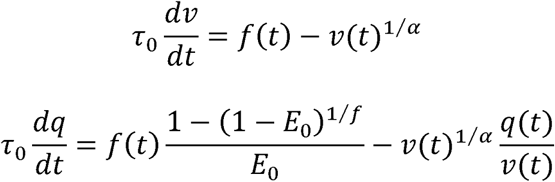

where *τ*_0_ is the hemodynamics transit time, *α* represents the resistance of the venous balloon (vessel stiffness), and *f*(*t*) is the blood inflow at time *t* and is given by

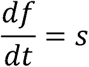

where *s* is an exponentially decaying, vasodilatory signal given by

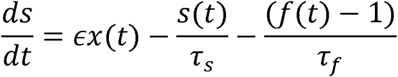

where *ϵ* is the efficacy with which neuronal activity *x*(*t*) (i.e., integrated synaptic activity) causes an increase in signal, *τ_s_*-is the time constant for signal decay, and *τ_f_*-is the time constant for autoregulatory feedback from blood flow (Friston et al., 2000). See Supplementary Table S6 for the values of the above parameters. The simulated fMRI BOLD time series resulting from the above equations were low-pass filtered (<0.25Hz) and down-sampled every two seconds.

### Resting State parameter exploration

We performed a global parameter exploration (for which we used exclusively the TVB simulator and the structural connectome with no task nodes) to obtain a reasonable match between empirical and model FC (Cabral et al., 2011). We obtained the empirical functional connectivity datasets from (Hagmann et al., 2008) which we used as a target for our simulated FC. Note that we used a low resolution (66 nodes) FC of matrices to perform the comparisons between empirical and resting state simulations (Honey et al., 2009): We transformed all correlation coefficients to Fisher’s Z values and averaged the FC matrices across subjects within each condition. We then calculated low-resolution (66 ROIs) matrices (each ROI corresponding to a brain region in the Desikan-Killiany parcellation (Desikan et al., 2006) for each condition (Hagmann et al., 2008; Honey et al., 2009) by averaging FC coefficients within each one of the low-resolution ROIs (Hagmann et al., 2008) and converted back to correlation coefficients using an inverse Fisher’s Z transformation. We systematically varied the global coupling parameter (*a*_Γ_ in the long-range coupling equation above) and the white matter conduction speed and conducted a 198-second resting state simulation for each parameter combination. We calculated a Pearson correlation coefficient between the model FC matrix (for each parameter combination) and the empirical FC matrix. Then, we chose the parameter combination that gave us the highest correlation value and used that combination for the PF, PV and DMS simulations of our study. The global strength parameter range used was between 0.0042 and 0.15 with a step of 0.01. The conduction speed parameter range used was between 1 and 10 m/s with a step of 1. The best combination of parameters was (0.15, 3) which yielded a correlation value between simulated and empirical FC of r = 0.37. Note that absent structural connections were removed from this correlation calculation as in (Honey et al., 2009), but not in the rest of the paper.

### From RS to PF, PV, and DMS

After finding an optimal match between empirical and simulated RS, we performed a simulation of RS with stimulation in visual task nodes using only the TVB simulator (Sanz-Leon et al., 2015). The correlation between RS FC and RS with stimulation FC was 0.90. Subsequently, we used a blend of our LSNM simulator and TVB to simulated PF. The correlation between RS with stimulation and PF was 0.9. As a last step, we performed a DMS simulation and compared it to the PF simulation (correlation was 0.79). Thus, we used a TVB RS simulation (matched to empirical RS) as a starting point for our PF and task-based simulations.

### Network construction

The simulations were performed using the TVB simulator with the 998-node Hagmann connectome and the LSNM visual short-term memory simulator described above. We isolated the synaptic activity timeseries of connectome nodes from the task nodes’ synaptic activity. We used the Balloon model to estimate fMRI BOLD activation over each one of the 998 nodes, for each condition, and for each subject separately. We calculated zero lag Pearson correlation coefficients for each pair of the BOLD timeseries to obtain a FC matrix for each condition and for each subject. We used the weighted FC matrices within each condition to construct graphs where each one of the 998 ROIs corresponded to a graph node and the correlation coefficients between each pair of ROIs corresponded to graph edges (Bolt, Nomi, et al., 2017; Di et al., 2013). To keep the same number of edges across conditions, we thresholded the network edges to a sparsity level of between 5% and 40% (Di et al., 2013) with a step size of 5%.

### Graph theory analysis

A set of eight graph theoretical metrics (global efficiency, local efficiency, clustering coefficient, characteristic path length, eigenvector centrality, betweenness centrality, participation coefficient, and modularity) were calculated using the FC matrices for each of the conditions using the Brain Connectivity Toolbox (Rubinov & Sporns, 2010) in Python, publicly available at https://github.com/aestrivex/bctpy. We calculated graph metrics for each individual FC matrix, for each condition and for each density threshold. Then we calculated the average and standard deviation of each graph metric for each density threshold.

*Global efficiency* (Latora & Marchiori, 2001) measures “functional integration” (Rubinov & Sporns, 2010) and indicates how well nodes are coupled through functional connections across the entire brain. Global efficiency is calculated as the average inverse shortest path length (Rubinov & Sporns, 2010). *Local efficiency* is the inverse of the average shortest path connecting a given node to its neighbors (Lee et al., 2017). *Clustering coefficient* (Watts & Strogatz, 1998) is a measure of “functional segregation” (Rubinov & Sporns, 2010). The clustering coefficient of a network node is the proportion of the given node’s neighbors that are functionally connected to each other. Whole brain clustering coefficient is calculated as the average of the clustering coefficients in a functional connectivity matrix (Rubinov & Sporns, 2010). *Characteristic path length* is the average shortest path length between all node pairs in a network (Rubinov & Sporns, 2010). *Eigenvector centrality* is a measure of centrality that considers degree of a given node and degree of that node’s neighbors (Fornito, Zalesky, & Bullmore, 2016 2016). Betweenness centrality is the fraction of shortest paths that cross a given network node (Rubinov & Sporns, 2010). *Participation coefficient* is a measure of each node’s participation in a given set of network communities. We used a set of six network communities for the participation coefficient calculation, as shown in Table S1 of (Hagmann et al., 2008), Table S1. *Modularity* (Newman, 2004) is a metric of functional segregation and it detects community structure in a network by dividing a functional connectivity matrix into sets of non763 overlapping modules and it measures how well a network can be divided into those modules (Rubinov & Sporns, 2010).

## SUPPORTING INFORMATION

**Table S1.**
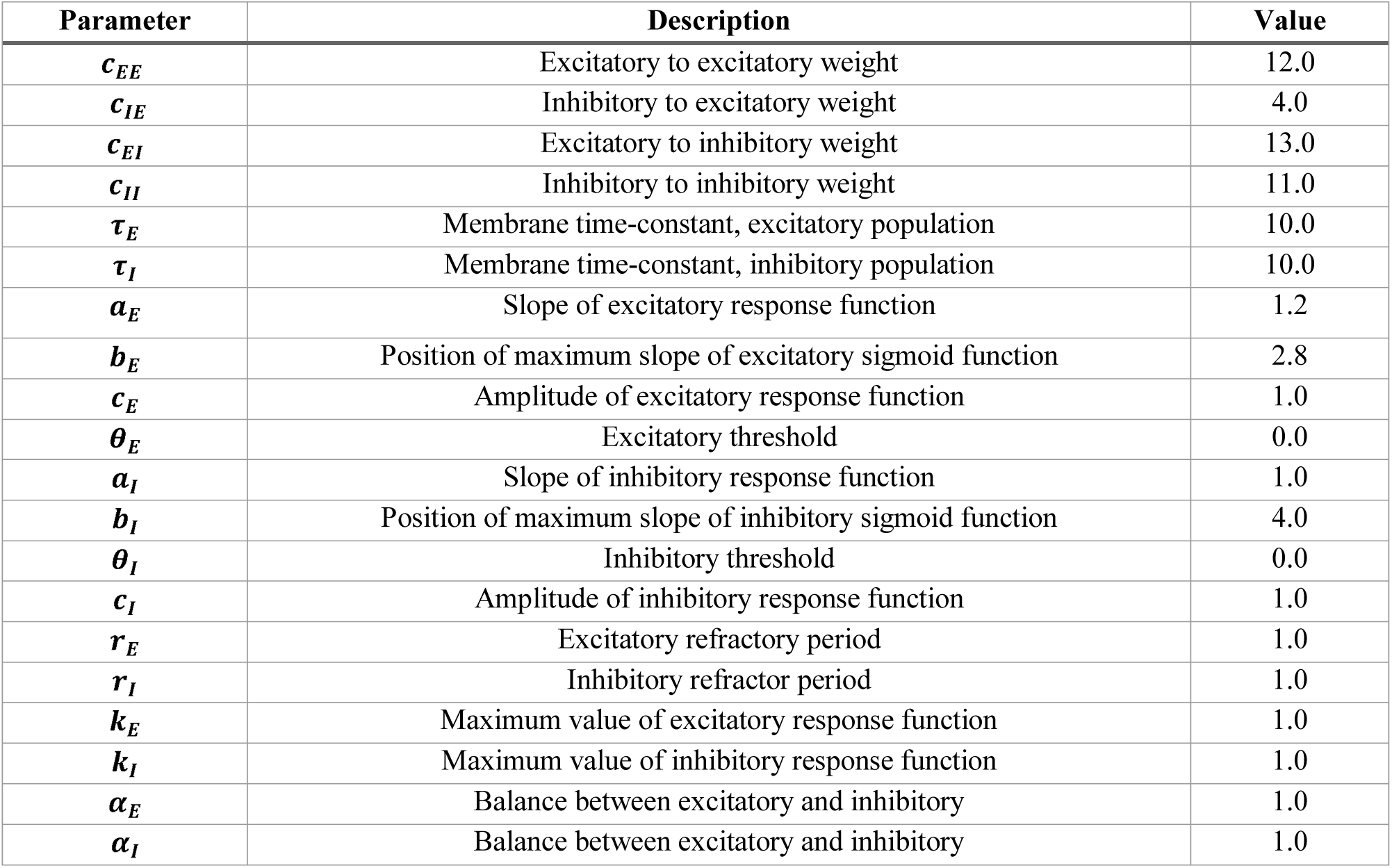
Parameters used in the Wilson-Cowan equation for each connectome node within TVB. The parameters shown above are the default parameters within TVB and are also shown in Table 11(a) of (Sanz-Leon et al., 2015).

**Table S2.** Parameters used for simulating the Hagmann et al. (Hagmann et al., 2008) connectome within the TVB resting state simulator. Please note the values of Global coupling strength and white matter transmission speed above are different to those presented in (Ulloa & Horwitz, 2016). In the present study we implemented a parameter search to better reproduce empirical RS FC of (Hagmann et al., 2008). See methods sections for details.

| Parameter | Value |
| --- | --- |
| Number of nodes | 998 |
| Global coupling strength | 0.15 |
| White matter transmission speed (mm/ms) | 3.0 |
| Integrator | Euler stochastic (dt=5) |

**Table S3.** Parameters used in the Wilson-Cowan unit model of each LSNM submodule

| Parameter | E element | I element |
| --- | --- | --- |
| $K$ | 9.0 | 20.0 |
| $\phi$ | 0.3 | 0.1 |
| $N$ | $\pm 0.025$ | $\pm 0.025$ |
| $\Delta$ | 0.5 | 0.5 |
| $\delta$ | 0.5 | 0.5 |

**Table S4.** Connection patterns among submodules of LSNM model

| Source | Destination | Fanout | Mean/SD | Percent to create | Comments |
| --- | --- | --- | --- | --- | --- |
| LGN | V1 | 7x7 | 34 @ 0.003±0.003<br>2 x 5 @ 0.006 ± 0.003<br>1 x 5 @ 0.020 ± 0.002 | 100 | Highest values oriented either vertically or horizontally |
| V1h | V4h | 1x5 | 0.04 ± 0.01 | 50 |  |
| V1v | V4v | 5x1 | 0.04 ± 0.01 | 50 |  |
| V1h | V4c | 3x3 | 4 @ 0.0 ± 0.01<br>5 @ 0.02 ± 0.01 | 50 | Lowest values at the corners |
| V1v | V4c | 3x3 | 4 @ 0.0 ± 0.01<br>5 @ 0.02 ± 0.01 | 50 | Lowest values at the corners |
| V4 | IT | 5x5 | 0.01 ± 0.01 | 50 | Learned |
| IT | FS | 1x1 | 0.2 ± 0.02 | 100 |  |
| D2 | V4 | 5x5 | 0.0014 ± 0.0007 | 100 |  |
| D1 | IT | 1x1 | 0.03 ± 0.001 | 100 | Inhibitory |
| D2 | IT | 1x1 | 0.01 ± 0.002 | 100 |  |
| IT | V4 | 4x4 | 0.00125 ± 0.0006 | 100 |  |

**Table S5.** Connection weights among submodules in the prefrontal cortex region of LSNM

| Source | Destination | Element | Weight |
| --- | --- | --- | --- |
| FS | D2 | E | 0.07 |
| FS | FR | E | 0.05 |
| D1 | FR | E | 0.06 |
| D1 | D2 | E | 0.105 |
| D2 | D1 | E | 0.10 |
| D1 | FS | I | 0.02 |
| FS | D1 | I | 0.05 |
| FR | D1 | I | 0.03 |
| FR | D2 | I | 0.065 |

**Table S6.**
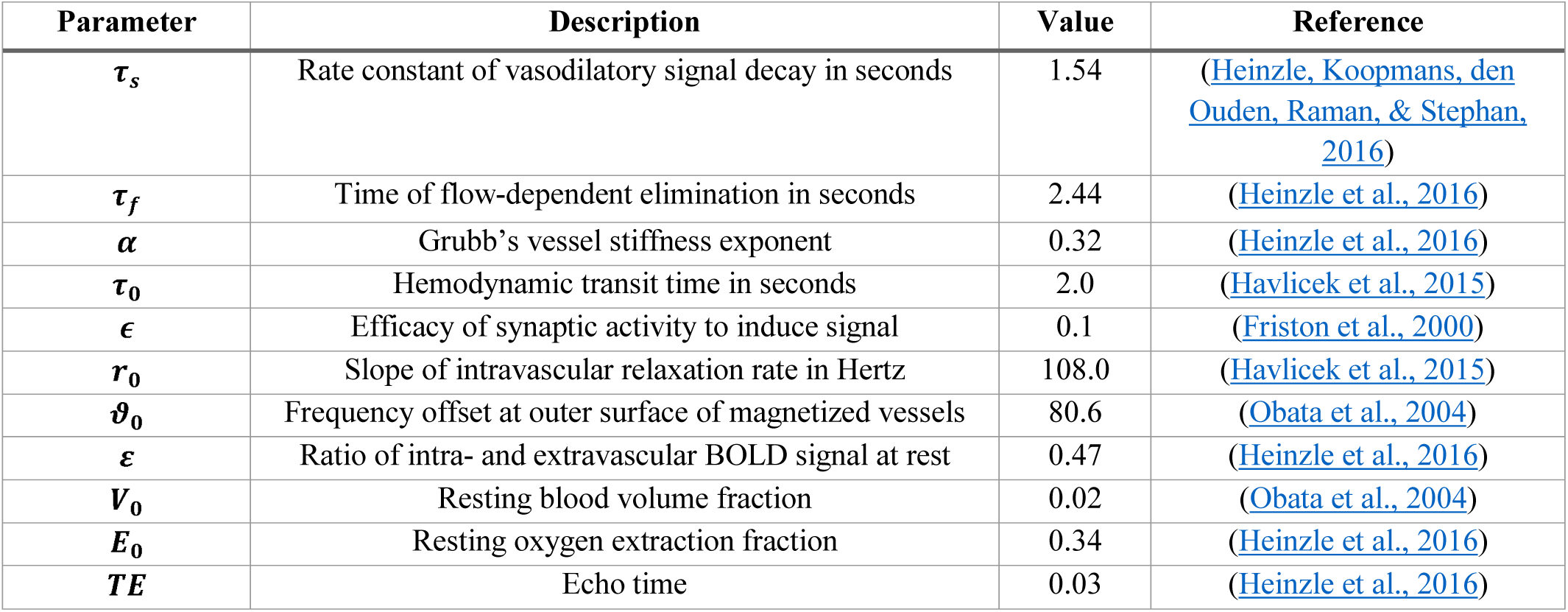
Parameters used for the Balloon model of hemodynamic response used in our simulations. Values are based on a 3T MRI magnet.

| Parameter | Description | Value | Reference |
| --- | --- | --- | --- |
| $\tau_s$ | Rate constant of vasodilatory signal decay in seconds | 1.54 | (Heinzle, Koopmans, den Ouden, Raman, & Stephan, 2016) |
| $\tau_f$ | Time of flow-dependent elimination in seconds | 2.44 | (Heinzle et al., 2016) |
| $\alpha$ | Grubb's vessel stiffness exponent | 0.32 | (Heinzle et al., 2016) |
| $\tau_0$ | Hemodynamic transit time in seconds | 2.0 | (Havlicek et al., 2015) |
| $\epsilon$ | Efficacy of synaptic activity to induce signal | 0.1 | (Friston et al., 2000) |
| $r_0$ | Slope of intravascular relaxation rate in Hertz | 108.0 | (Havlicek et al., 2015) |
| $\theta_0$ | Frequency offset at outer surface of magnetized vessels | 80.6 | (Obata et al., 2004) |
| $\varepsilon$ | Ratio of intra- and extravascular BOLD signal at rest | 0.47 | (Heinzle et al., 2016) |
| $V_0$ | Resting blood volume fraction | 0.02 | (Obata et al., 2004) |
| $E_0$ | Resting oxygen extraction fraction | 0.34 | (Heinzle et al., 2016) |
| $TE$ | Echo time | 0.03 | (Heinzle et al., 2016) |

## ACKNOWLEDGEMENTS

This research was funded by the Division of Intramural Research of the National Institute on Deafness and Other Communication Disorders. We thank Olaf Sporns and Chris Honey for sharing the functional and structural connectivity data sets from their empirical studies used in the present paper. We thank Paul Corbitt for useful discussions related to the simulation code used for our analysis and the parameters used for converting synaptic activity to fMRI BOLD time-series. We thank Marmaduke Woodman for helping us navigate technical aspects of the TVB simulator.

